# VopU is a novel T3SS effector protein with mono-ADP-ribosyltransferase activity and a non-canonical H-Y-Q catalytic triad

**DOI:** 10.64898/2026.05.07.723553

**Authors:** Sebastián A Jerez, María J Puentes-Maldonado, Víctor Cabrera, Fernando A Amaya, Linda J. Kenney, Carlos A Santiviago, Carlos J Blondel

## Abstract

*Vibrio parahaemolyticus* utilizes its second Type III Secretion System (T3SS2) to deliver a suite of effector proteins that subvert eukaryotic host cell processes. In this study, we characterize VPA1312, renamed VopU, as a novel T3SS2 effector. We demonstrate that VopU is translocated into infected cells via the T3SS2 and independently of the VocC chaperone. From bioinformatic analyses, VopU is a member of a family of proteins distributed across diverse bacterial taxa and in some cases associated with T6SS gene clusters and plasmids. Sequence, structure-based, and functional analysis reveals that VopU harbors an ADP-ribosyltransferase domain with a non-canonical H-Y-Q catalytic triad, a motif not previously described in naturally occurring mono-ADP-ribosyltransferases. Heterologous expression of VopU within transfected cells or translocation of VopU during infection results in the ADP-ribosylation of a small protein of approximately 18 kDa protein distinct from the target of the previously described ADP-ribosyltransferase effector, VopT. While VopU is not essential for T3SS2-dependent cytotoxicity or intracellular survival of *V. parahaemolyticus*, heterologous expression of VopU induces host cell rounding and a transcriptional host stress response, likely linked to NAD depletion. These findings provide the initial characterization of this novel family of mono-ADP-ribosyltransferases and expand the known repertoire of bacterial effectors that mediate orthogonal post-translational modifications during host infection.

**IMPORTANCE:** *Vibrio parahaemolyticus* is a leading cause of seafood-borne illness worldwide. To cause disease, the bacterium relies on a Type III Secretion System to deliver effector proteins into the cytosol of infected cells, subverting cellular processes. In this study, we identified a novel effector protein, VopU, which performs a post-translational modification known as ADP-ribosylation using a non-canonical catalytic triad (H-Y-Q). This is the first description of a naturally occurring mono-ART with this specific motif. We showed that VopU homologs are distributed across other bacterial species, and that VopU is active during infection of host cells. Together, our findings identify a novel family of effectors that can modify host proteins and potentially contribute to host colonization during infection.

## INTRODUCTION

*Vibrio parahaemolyticus* is a Gram-negative marine bacterium that is the leading cause of seafood borne gastroenteritis worldwide (1). In 1996, a new clonal strain of the O3:K6 serotype emerged that has been responsible for the major global outbreaks of *V. parahaemolyticus* (2–4). The pathogenic potential of the pandemic clone and its derivatives is mostly attributed to the Type III Secretion System 2 (T3SS2), which is encoded within an 80 kb genomic island located in chromosome II known as the *V.* parahaemolyticus pathogenicity island 7 (VpAI-7) (5). The T3SS2 is essential for intestinal colonization, fluid accumulation, and the inflammatory diarrhea observed during infection with *V. parahaemolyticus* (6–9), and requires host cell surface fucosylation to cause cytotoxicity (10). Notably, the T3SS2 gene cluster is not restricted to pandemic *V. parahaemolyticus* strains, as it extends beyond the *Vibrionaceae* family (11).

T3SSs are complex nanomachines that enable Gram-negative bacteria to deliver proteins known as effectors directly from the bacterial cytosol into the cytosol of eukaryotic cells. Translocation of effectors into host cells enables pathogens to subvert a wide range of cellular functions (12–15). In this context, the contribution of T3SSs to the pathogenic potential of human, animal and plant pathogens depends on the repertoire of effector proteins translocated into host cells and on the intracellular networks generated during infection (15–18).

A key strategy employed by bacterial pathogens to subvert host biology is the delivery of effector proteins that can modify host proteins post-translationally (13, 19–21). In this context, a wide range of bacterial pathogens deliver toxins and effector proteins that catalyze the transfer of an ADP-ribose group from NAD+ to specific eukaryotic target molecules, in a process known as ADP-ribosylation, which can drastically alter protein function (22–25). Bacterial ADP-ribosyltransferase toxins (bARTTs) are typically classified into clades based on conserved catalytic residues, such as the H-Y-[EDQ] (Diphtheria toxin-like) and the R-S-E (cholera toxin-like) families (22, 23). Members of the H-Y-[EDQ] clade, such as Diphtheria toxin (DT) and *Pseudomonas* Exotoxin A (ExoA), utilize a canonical H-Y-E (Histidine-Tyrosine-Glutamate) catalytic triad to modify elongation factor 2 (eEF2), thereby inhibiting protein synthesis and inducing cell death (26, 27). Members of the R-S-E family have a more diverse range of host targets, including actin, ubiquitin and G proteins (23). Interestingly, there are increasing reports of novel bARTTs delivered by specialized protein secretion systems such as the T3SS and Type VI Secretion systems (T6SS) (28–34).

To date, twelve T3SS2 effector proteins have been identified within the T3SS2 gene cluster of *V. parahaemolyticus* strain RIMD2210633 (reviewed in (1, 35)). However, the full repertoire of effector proteins remains unknown. Of the repertoire of T3SS2 effector proteins described to date, only VopT has been shown to correspond to a bARTTs (36). VopT is an ADP-ribosylstransferase of the R-S-E family which contributes to T3SS2-dependent cytotoxicity of *V. parahaemolyticus* through the ADP-ribosylation of the host factor G protein Ras (36). Notably, there are no reports of additional bARTTs delivered by the T3SS2 of *V. parahaemolyticus*.

Recent reports have suggested that the uncharacterized VPA1312 open reading frame (ORF) is a component of the T3SS2 of *V. parahaemolyticus* (37–39). Earlier work has shown that VPA1312 is expressed under T3SS2-inducing conditions (39) and that the VPA1312 protein interacts with the gatekeeper protein VgpA (38), even though it is not required for T3SS2-induced cell death during infection (37).

In this study, we provide evidence that VPA1312, now VopU, is a novel T3SS2 effector protein with mono-ADP-ribosyltransferase (mono-ART) activity. VopU is translocated into infected cells in a T3SS2-dependent manner, but independently of the previously described VocC chaperone. Bioinformatic analyses revealed that VopU was part of a wider family of proteins with homologs in other bacterial species, including *E. coli* and *Bordetella*. Interestingly, we show that VopU harbors a non-canonical H-Y-Q catalytic triad, which was required for ADP-ribosylation of a small protein of approximately 18 kDa when VopU was heterologous expressed in eukaryotic cells and during infection of HeLa cells by *V. parahaemolyticus*. In agreement with a previous report (37), VopU does not contribute to T3SS2-dependent cell death, even though heterologous expression experiments suggest it is involved in changes in cell morphology and NAD depletion when expressed. We also provide evidence that VopU and VopT are the major contributors to mono-ADP-ribosylation of host proteins during infection by *V. parahaemolyticus*.

## RESULTS

### VPA1312 encodes a novel T3SS2 effector (VopU) that is translocated independently of VocC

Bioinformatic analyses of previously uncharacterized ORFs within Vpai-7 of *V. parahaemolyticus* identified VPA1312. This ORF encodes a 174-amino acid protein with an N-terminal region (residues 1-99) that shares similarity with the Diphtheria toxin domain (CATH superfamily 3.90.175.10) (**Figure 1A**). Previous studies have demonstrated that VPA1312 is expressed under T3SS2-inducing conditions and secreted via the T3SS2 apparatus (37, 39). The presence of a putative toxin domain and its T3SS2-dependent secretion (37) suggested that VPA1312 was likely a T3SS2 effector protein.

**Figure 1.**
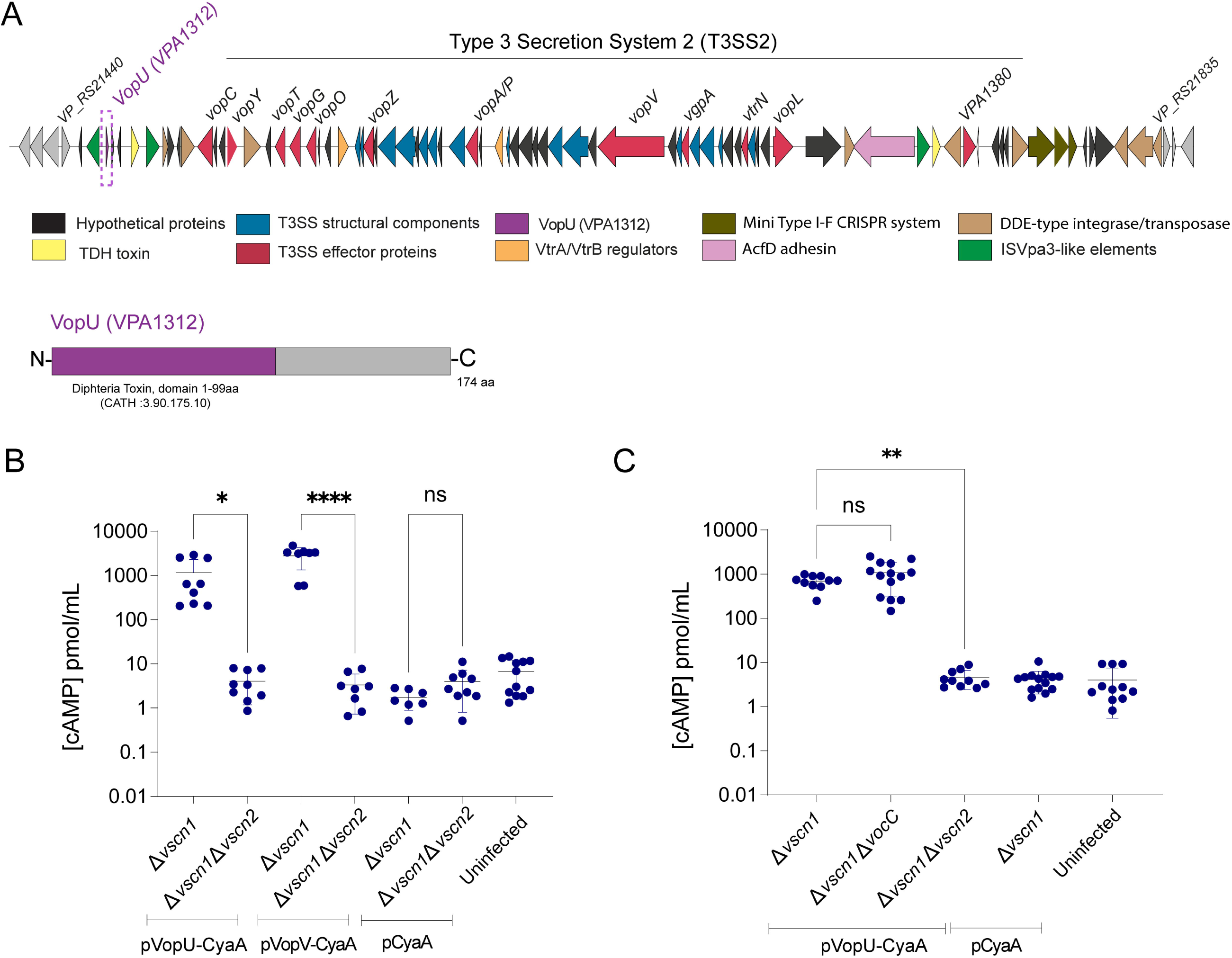
VPA1312 (VopU) is T3SS2 effector protein. **(A)** Schematic representation of the genetic organization of the T3SS2 gene cluster in *V. parahaemolyticus* RIMD2210633 and the predicted ART domain in VopU. Translocation of VopU-CyaA and VopV-CyaA was assessed via determination of intracellular cAMP levels (pmol/ml) in infected Caco-2 cells **(B)** and HT-29 cells **(C)**. Values are means plus standard deviations (error bars) from at least two independent biological replicates each with three technical replicates. Asterisks indicate significant differences (*t* test, ***, *P* < 0.001). ns, not significantly different.

To determine whether VPA1312 corresponded to a bona fide effector protein, a gene fusion was constructed between VPA1312 and the *cyaA* gene from *Bordetella pertussis* in the plasmid pCyaA. Translocation experiments performed in both Caco-2 and HT-29 cells showed T3SS2-dependent translocation of VPA1312-CyaA, revealed by an increase in intracellular cyclic adenosine monophosphate (cAMP) levels, measured by enzyme-linked immunosorbent assay (ELISA) (**Figure 1B and 1C**). A VopV-CyaA reporter served as a positive control for T3SS2-dependent translocation. As shown in **Figure 1B**, infection of Caco-2 cells with *V. parahaemolyticus* strains expressing VPA1312-CyaA resulted in a significant, T3SS2-dependent increase in cAMP levels, comparable to the VopV positive control. The same T3SS2-dependent translocation was observed when HT-29 were used cells (**Figure 1C**). Effector proteins typically require chaperone proteins for proper folding and delivery to the T3SS sorting platform (40–43). To date, only one chaperone, VocC, has been described for T3SS2 (44). To investigate the mechanism of VPA1312 translocation, experiments were carried out with a *V. parahaemolyticus* strain lacking *vocC*. As shown in **Figure 1C**, the absence of VocC did not affect the ability of *V. parahaemolyticus* to translocate VPA1312 in a T3SS2-dependent manner, suggesting the involvement of an as-yet uncharacterized chaperone or a chaperone-independent translocation. Collectively, these data indicate that VPA1312 encodes a novel T3SS2 effector protein. Accordingly, we propose to name this effector VopU, in accordance with the nomenclature currently used for T3SS2 effector proteins in *Vibrio* (35).

### VopU shares similarity to bacterial ARTs of the H-Y-[EDQ] clade

As mentioned above, bioinformatic analysis suggested the presence of a putative ADP-ribosyltransferase (ART) domain at the N-terminal region of VopU (**Figure 1A**). To gain further insight into this putative domain, we performed additional sequence- and structure-based analyses of the VopU amino acid sequence. Sequence-based analysis showed that VopU shares similarity with the N-terminal region of ART domains of bacterial toxins, such as ExoA of *Pseudomonas aeruginosa*, Diphtheria toxin of *Corynebacterium diphtheriae*, and Cholix toxin of *Vibrio cholerae* (**Figure 2A**). Each of these toxins corresponds to bARTTs of the H-Y-[EDQ] family, with a catalytic triad consisting of a histidine, tyrosine, and glutamate (H-Y-E). The analysis showed conservation of the catalytic histidine and tyrosine residues (H7 and Y39 in VopU) within the N-terminal region of VopU, with a more divergent C-terminal region (**Figure 2A**).

**Figure 2.**
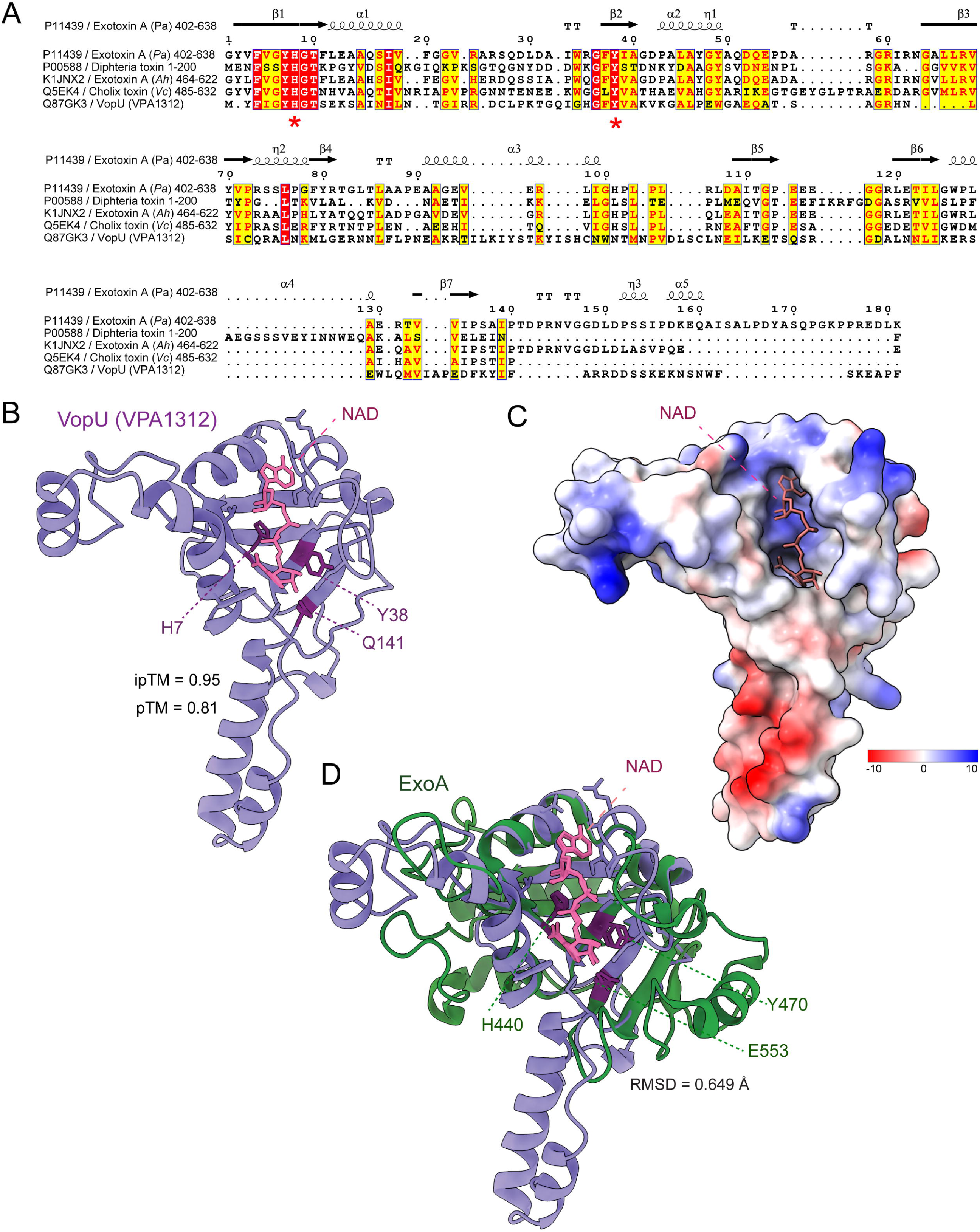
VopU is similar to bacterial H-Y-E ARTs. **(A)** Multiple sequence alignment of VopU and the ART domain of bacterial toxins of H-Y-[EDQ] family. BLASTp alignments were performed using T-Coffee Expresso and visualized by ESPript 3.0. Amino acids with a red background correspond to positions with 100% identity; amino acids with a yellow background correspond to positions with >70% identity. UniProt IDs and amino acid region analyzed of each protein are shown at the beginning of each line of the alignment. (**B**) Alphafold3 predicted structure of VopU in complex with NAD. The predicted catalytic H-Y-Q triad is highlighted in purple and iPTM and pTM scores are shown. (C) Molecular surface and electrostatic potential of the Alphafold3 predicted structure of VopU in complex with NAD. (**D**) Superimposition of the predicted structure of VopU with the crystalized structure of ExoA in complex with NAD (PDB 2ZIT) highlighting the H-Y-E catalytic triad of ExoA superimposed to the H-Y-Q triad of VopU.

Analysis of the Alphafold3-predicted structure of VopU identified a putative NAD-binding pocket and the presence of an H-Y-Q catalytic triad (H7, Y39, and Q141) in complex with NAD+ (**Figure 2B and 2C**). Thus, while the identification of the catalytic histidine and tyrosine are in agreement with sequence alignment analyses (**Figure 2A**), the structural analysis allowed the identification of the putative catalytic glutamine residue within the more divergent C-terminal region of VopU. Analysis of the predicted structure of VopU superimposed on the structure of ExoA further supported the presence of a conserved NAD binding pocket (**Figure 2D**). In addition, the ExoA catalytic triad H-Y-E overlapped with the non-canonical H-Y-Q triad predicted for VopU (**Figure 2D**).

To the best of our knowledge, no previously identified bARTTs have been described as possessing a H-Y-Q catalytic triad. While the binding pocket of ExoA and VopU was structurally conserved, there was little overlap in the rest of the predicted protein structure (**Figure 2D**). These structural differences and the presence of a non-canonical catalytic triad (H-Y-Q versus H-Y-E), suggest that VopU might act on different host substrates compared to previously described bARTTs of the H-Y-[EDQ] family.

### VopU is predominantly distributed within *Vibrio* species, although it is also detected in other bacterial taxa

It has been reported that the T3SS2 gene cluster is not restricted to the *Vibrionaceae* family and that T3SS2 gene clusters can differ in the distribution of effector proteins encoded within them (11). In this context, we analyzed the presence and distribution of VopU homologs in bacterial genomes. Sequence-based BLASTp analyses identified 2007 homologs distributed across 9 bacterial genera, including *Vibrio*, *Escherichia*, *Bordetella*, and *Shewanella*, among others (**Figure 3A and 3B**). Despite this distribution, most VopU homologs were found in *Vibrio* species, and most (92%) in *V. parahaemolyticus* strains (**Figure 3A**).

**Figure 3.**
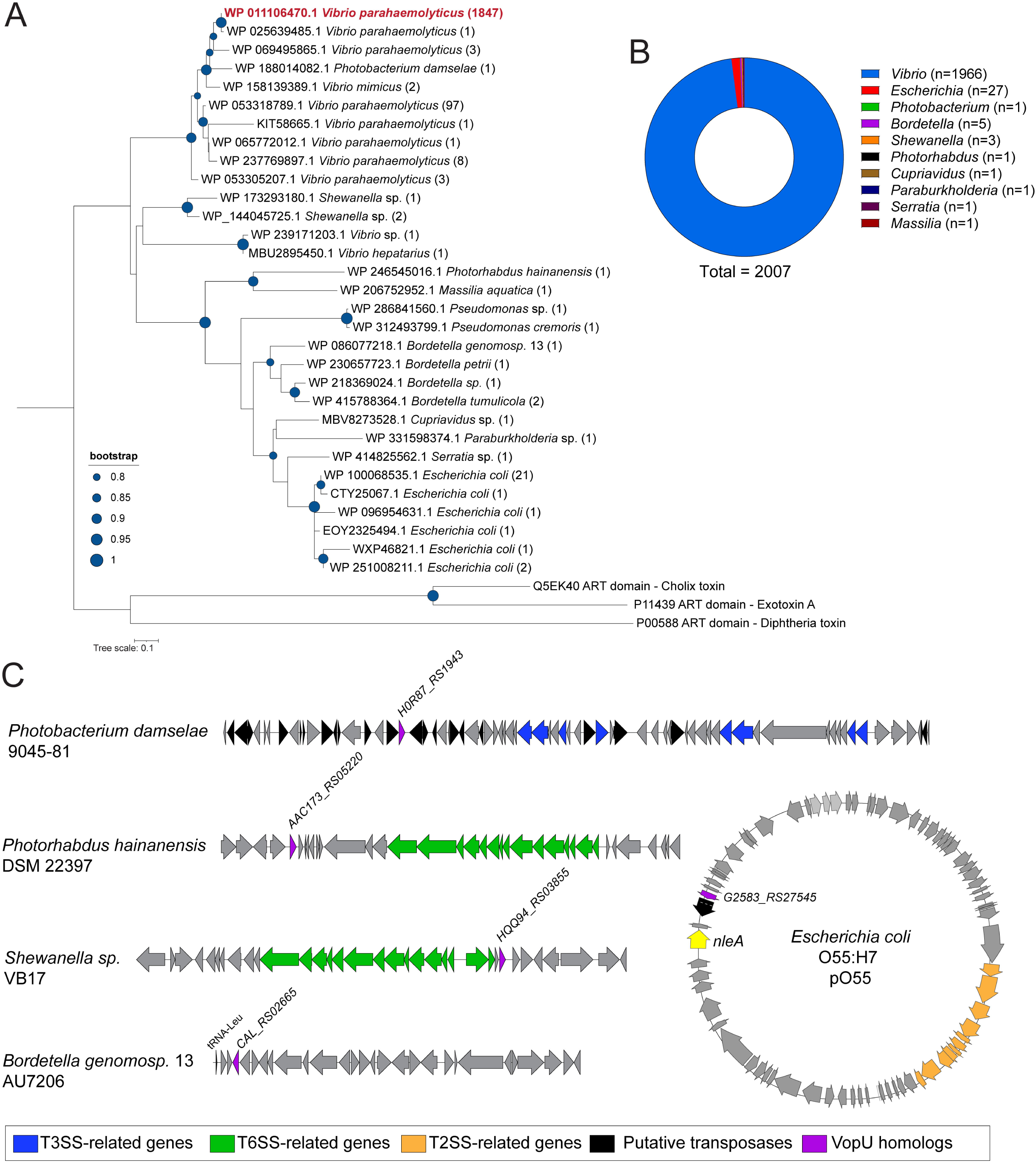
VopU homologs are mostly distributed among *Vibrio, Bordetella* and *E. coli*. **(A)** Phylogenetic analysis of the VopU homologs identified in this study. Phylogenetic analysis was performed with MEGA and visualized by iTOL. VopU homologs in *V. parahaemolyticus* genomes is highlighted in red color. **(B)** Distribution of VopU homologs in terms of bacterial genera. **(C)** Schematic depiction of the genetic context of VopU homologs in different bacterial genomes.

Phylogenetic analyses showed that VopU homologs form a distinct cluster distinguishable from other related ARTs such as Diphtheria, Exotoxin A, and Cholix toxin. In addition, phylogenetic analysis revealed differences among VopU homologs, with those identified in *E. coli*, *Bordetella*, and *Pseudomonas* clustering into distinct internal branches of the VopU family (**Figure 3A**). A more in-depth analysis of the genetic context of genes encoding VopU homologs found that, while in *Vibrio* species they are located within or near T3SS2 gene clusters, in other bacterial species they are found at more diverse locations (**Figure 3C**). For example, in *Photorhabdus* and *Shewanella* species, genes encoding VopU homologs were often found close to T6SS gene clusters. In contrast, in *E. coli*, VopU homologs were always encoded on plasmids related to the pO55 plasmid of *E. coli* O55:H7, which also encodes the NleA T3SS effector protein (**Figure 3C**).

Structure-based analysis also identified VopU as a member of a larger family of proteins. The Foldseek clustered AlphaFold database, AFDB Clusters (45), identified VopU as a member of cluster A0A1W6Z7P8. Members of this cluster include predicted structures from some of the *E. coli*, *Bordetella* and *Shewanella* homologs identified by the sequence-based analyses mentioned above. In summary, these results suggest that VopU is the founding member of a new H-Y-Q bARTTs class distributed beyond the *Vibrio* genus.

### VopU does not contribute to T3SS2-dependent cytotoxicity or intracellular survival of *V. parahaemolyticus*

The T3SS2 delivers over a dozen effector proteins that contribute to the adhesion, invasion, proliferation, and cytotoxicity of infected host cells (reviewed in (1, 35)). To determine whether VopU contributes to these processes, we constructed a *V. parahaemolyticus* mutant strain harboring a *vopU* deletion in the Δ*vscn1* genetic background and performed infection experiments in HT-29 and Caco-2 intestinal epithelial cells.

To evaluate cellular toxicity, we used two common methods that rely on the disruption of the cellular membrane: i) a fluorescent DNA-binding dye that enters the cell upon cellular damage, and ii) the determination of cellular release of the LDH enzyme to the supernatant of infected cells. As previously reported, a *V. parahaemolyticus* strain harboring a functional T3SS2 (Δ*vscn1*, T3SS2+) caused time-dependent cytotoxicity not observed when a *V. parahaemolyticus* strain lacking both T3SSs was employed (Δ*vscn1* Δ*vscn2*) (**Figure 4A and 4B**). In contrast, infection with the strain lacking *vopU* did not show significant differences in comparison to the T3SS2+ strain (**Figure 4A and 4B**). This result indicated that VopU was not a major contributor to T3SS2-dependent host cell death.

**Figure 4.**
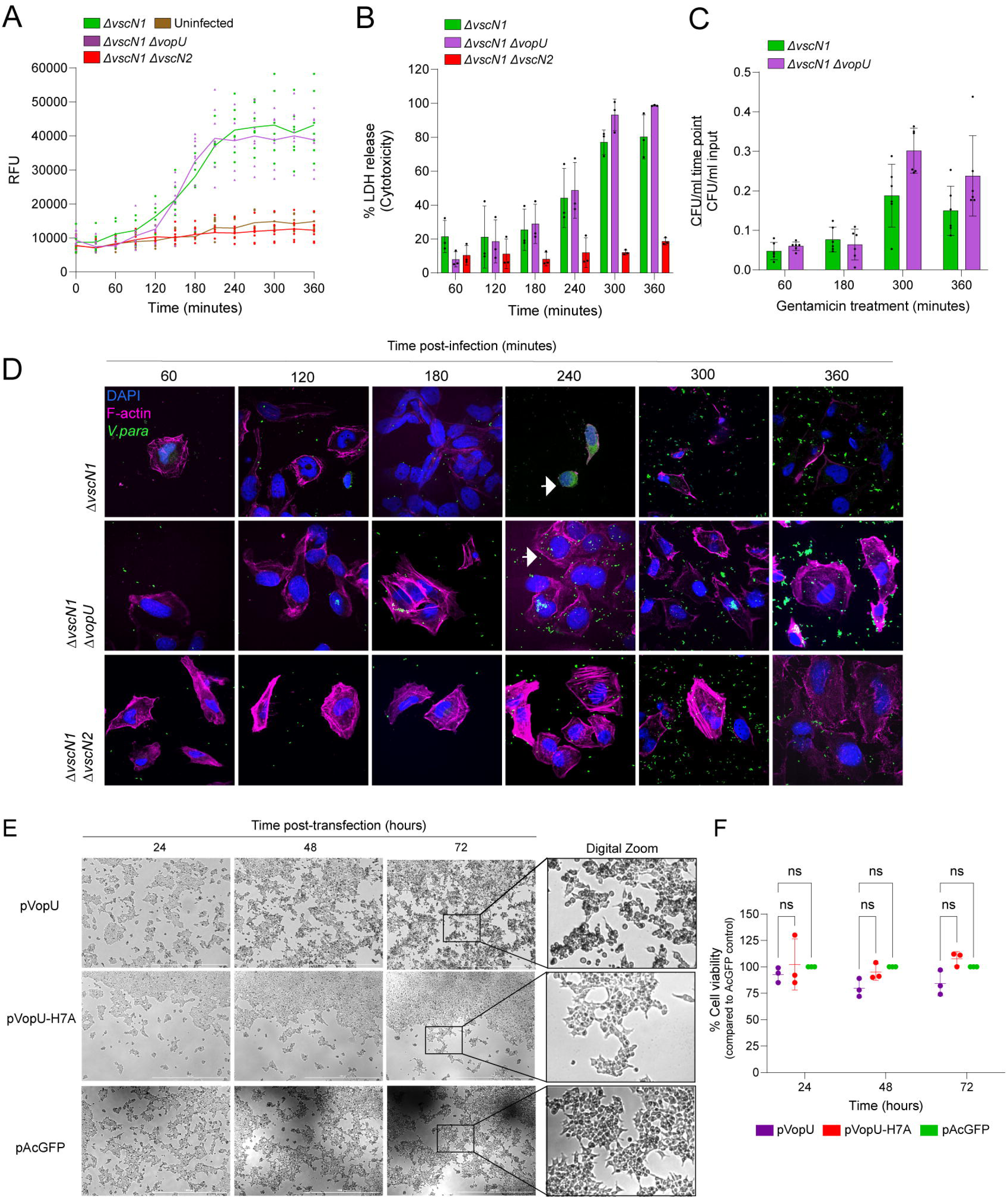
VopU is not a major contributor to T3SS2-dependent cytotoxicity, invasion or intracellular survival of *V. parahaemolyticus*. Caco-2 BBE cells were infected with different strains of *V. parahaemolyticus* and cellular toxicity was evaluated by CellTox Green Cytotoxicity assay **(A)** and LDH release analysis **(B)** Values are means from at least two independent biological replicates each with three technical replicates. **(C)** Gentamicin protection assay of cells infected with *V. parahaemolyticus*. **(D)** Confocal microscopy analysis of HeLa cells infected with *V. parahaemolyticus* stains harboring the pGFP plasmid. F-actin and nuclear DNA were visualized by phalloidin and DAPI staining. White arrows point to cells with different cell shape. **(E)** Heterologous expression of VopU and a VopU variant with a mutation in the predicted catalytic histidine (H7A) in HEK293T cells. Overexpression of AcGFP was used as a control. Digital zoom highlights differences in cell shape. **(F)** Cell viability of HEK293T cells which express each VopU, VopU H7A and AcGFP. Viability was assessed by trypan blue exclusion at different time points. ns, not significantly different (*t* test).

In addition to causing host cell death, *V. parahaemolyticus* can invade and proliferate within infected cells (46–49). To test whether VopU could contribute to these processes, we assessed invasion and intracellular survival of the *vopU* deletion mutant strains in HeLa cells by means of a gentamicin protection assay. No differences in invasion or intracellular survival were observed between the *vopU* deletion mutant strain and the parental Δ*vscn1* strain, even though at later time points of infection, there was a trend toward higher intracellular colony-forming units (CFUs) in the strain lacking *vopU*, albeit not statistically significant (**Figure 4C**). Confocal microscopy analysis of HeLa cells infected with *V. parahaemolyticus* strains constitutively expressing the green fluorescent protein (GFP) showed that *V. parahemolyticus* caused changes in cellular morphology (evaluated by partial loss of phalloidin actin staining) at later time points as previously described (46) (**Figure 4D**). Interestingly, infection of cells with the *vopU* deletion mutant strain did not show the same changes in cellular morphology (**Figure 4D**).

To avoid potential redundancy with other T3SS2 effectors, we decided to express VopU and a derivative in which the predicted catalytic histidine was replaced with an alanine (H7A) by transfecting HEK293T cells with a eukaryotic expression vector that expresses each protein under a strong CMV promoter. As a control, a vector expressing *Aequorea coerulescens* GFP (AcGFP) under the control of the same promoter was used. Analysis of transfected cells over time (**Figure 4E and 4F**) showed that after 48-72 hours post-transfection, cells that expressed VopU exhibited a round morphology, which was not observed when cells expressed the H7A variant or the AcGFP control (**Figure 4E**). In addition, no changes in cellular viability were observed by either expressing VopU or the H7A variant in comparison to the AcGFP-transfected control cells (**Figure 4F**). These results suggest that expression of VopU within cells, but not a catalytic deficient variant, induced cellular stress, leading to changes in cellular morphology, but not cell death, at least during the time points examined.

### VopU-mediated ADP-ribosylation activity requires a H-Y-Q catalytic triad

Attempts to purify VopU were unsuccessful due to its tendency to form inclusion bodies, so we decided to use the heterologous expression approach outlined above to determine its predicted ART activity and assess the contribution of the predicted H-Y-Q catalytic triad to this process. First, we transfected HeLa cells for 48 hours with plasmids expressing VopU, the VopU H7A variant, and the AcGFP control, and analyzed the ADP-ribosylated protein profile from whole-cell lysates. To detect ADP-ribosylated proteins, we used a recently described highly specific antibody for mono-ADPr (AbD43647) (50, 51). As shown in **Figure 5A**, transfection of VopU led to the identification of an ADP-ribosylated protein of approximately 18 kDa, which was not detected when cells were transfected with the AcGFP control or when exposed to hydrogen peroxide as an inducer of host ADP-ribosylation upon cellular stress (**Figure 5A**). Notably, the identified protein was not detected when VopU-H7A was used, suggesting that histidine 7 is important for the ART activity of VopU.

**Figure 5.**
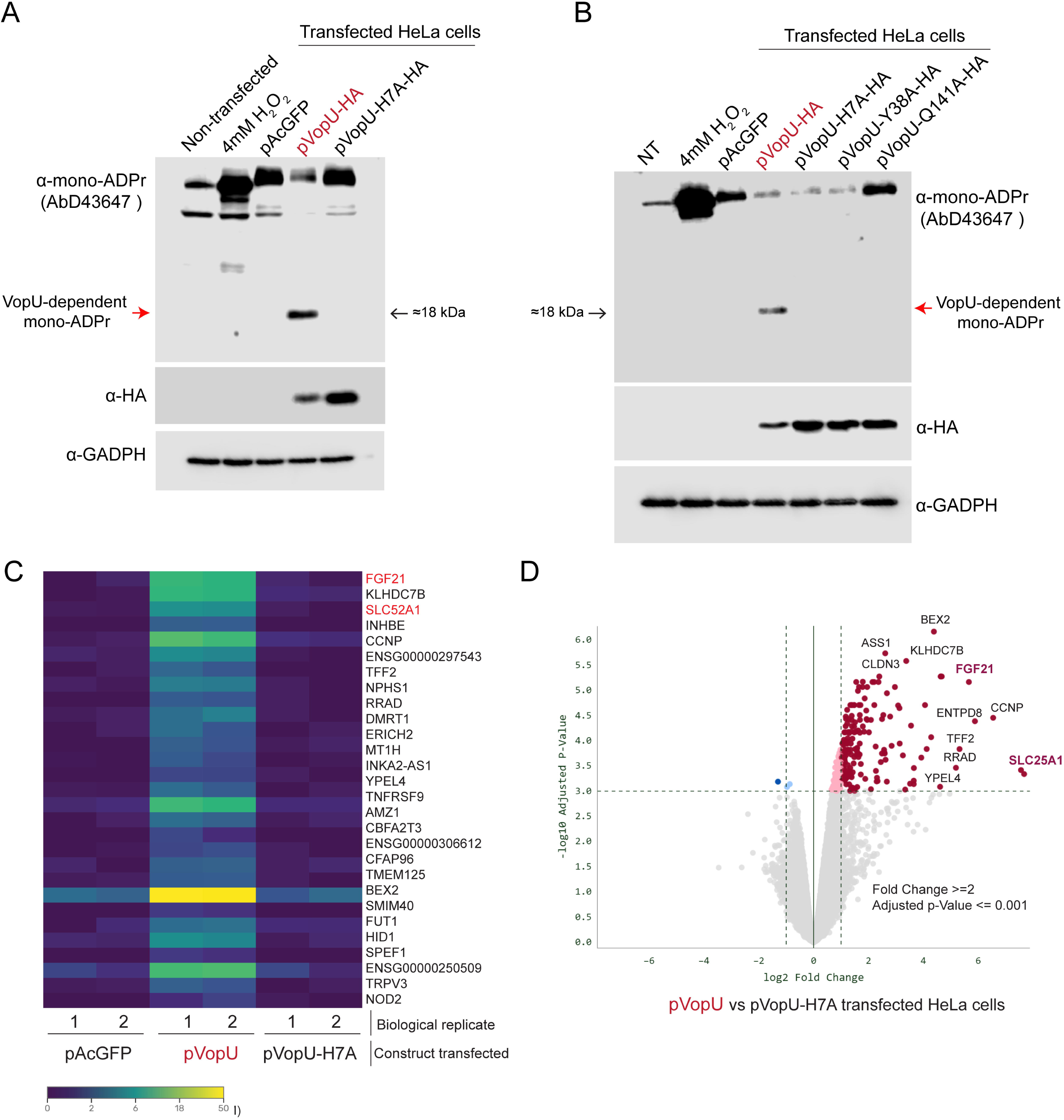
VopU is a mono-ART with a non-canonical H-Y-Q catalytic triad. SDS/PAGE and Western blot analysis of the profile of ADP-ribosylated proteins in HeLa cells. Cells were either treated with hydrogen peroxide to cause stress-induced ADP-ribosylation or transfected for 48 hours with constructs expressing either VopU, VopU-H7A **(A)** or VopU-Y38A and VopU-Q141A **(B)** expressing an HA epitope.ADP-ribosylation was analyzed by a mono-ADPr antibody (AbD43647). Production of VopU and H7A, Y38A and Q141 variants were analyzed with an antibody against the HA tag. Detection of host cellular GADPH was used as a loading control. Expression of AcGFP was used as a control. **(C)** RNAseq analysis of HeLa cells transfected with either VopU, VopU-H7A or AcGFP. Cells were collected after 48 hours of transfection from two biological replicates. The heatmap of gene read counts per million (CPM) shows the profile of the 28 locus with the greatest change in gene expression. **(D)** Volcano plot of the RNAseq analysis highlighting genes with greater than 2-fold change in gene expression and an adjusted p-Value of less than 0.001.

To test if the remaining amino acids conforming the predicted catalytic triad of VopU contributed to ADP-ribosylation, we generated VopU variants where residues Y38 and Q141 were substituted with alanine (Y38A and Q141A). As shown in **Figure 5B**, the ≈18 kDa protein ADP-ribosylated by VopU was not detected when the H7A, Y38A, and Q141A variants were expressed in HeLa cells. These results suggest that VopU is a functional mono-ART with an H-Y-Q catalytic triad.

To gain insights into the consequences of VopU expression, we determined the transcriptional profile of transfected cells. HeLa cells were transfected with constructs expressing VopU, the VopU H7A variant, and AcGFP as a control. After 48 hours post-transfection, cells were harvested and analyzed by RNAseq. Analysis of the transcriptome of transfected cells by RNAseq analyses, revealed that cells transfected with the AcGFP control had a transcriptional profile similar to that of cells expressing the inactive VopU-H7A variant, but was significantly distinct from that of cells transfected with a functional VopU (**Figure 5C**). In addition, most genes upregulated in cells expressing VopU were involved in metabolic cellular stress processes, some of which were directly linked to NAD depletion (e.g., SLC25A1 and FGF21) (**Figure 5D**), further supporting the notion that VopU is a functional bARTTs.

### VopU and VopT mono-ADP-ribosylate distinct host proteins during infection

To date, only one other T3SS2 effector protein with ADP-ribosylation activity has been identified, namely VopT, which is responsible for ADP-ribosylation of the Ras protein (36). The experiments highlighted above indicated that VopU expression led to ADP-ribosylation of a protein of approximately 18 kDa. To compare ribosylation profiles, we infected HeLa cells with *V. parahaemolyticus* strain derivatives of the Δ*vscn1* strain (T3SS2+) lacking either VopU or VopT, and we determined the profile of ADP-ribosylated proteins during infection. As shown in **Figure 6A**, infection with the parental Δ*vscn1* (T3SS2+) strain generated a profile composed of at least 5 different mono-ADP-ribosylated proteins, including a protein of approximately 18 kDa. Notably, infection with a Δ*vscn1* derivative lacking VopU generated a similar profile of ADP-ribosylated proteins, with the sole exemption of the 18 kDa protein, suggesting that this protein was ADP-ribosylated by VopU. Furthermore, infection with a Δ*vscn1* derivative lacking VopT led to the identification of only the 18 kDa protein. Taken together, these results suggest that VopU was responsible for the specific ADP-ribosylation of an approximately 18 kDa protein, while VopT mediates the ADP-ribosylation of at least four other proteins (in addition to Ras) during infection (**Figure 4B**).

**Figure 6.**
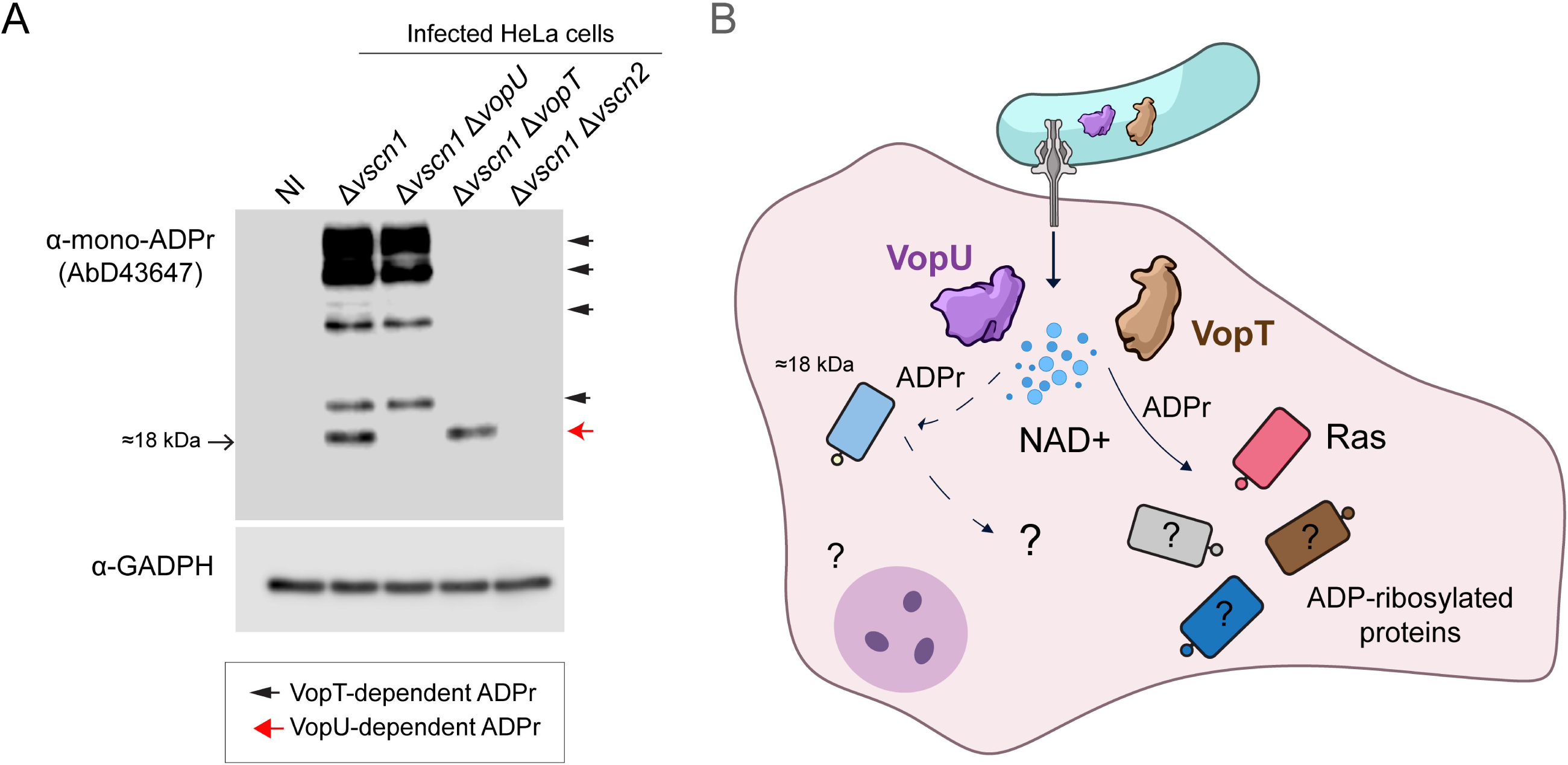
VopU and VopT generate a distinct mono-ADP-ribosylation profile during HeLa infection by *V. parahaemolyticus*. **(A)** HeLa cells were infected for 3 hours with *V. parahaemolyticus* strains and the profile of mono-ADP-ribosylated proteins was analyzed by SDS/PAGE and Western Blot analyzes. Uninfected cells and cells infected with a *V. parahaemolyticus* strain lacking functional T3SSs were included as controls. Different color and size arrows highlight the profile of VopU and VopT-dependent ADP ribosylations. Detection of host GADPH was used as a loading control. **(B)** Proposed model of cellular mono-ADP-ribosylation caused by VopU and VopT during host infection with *V. parahaemolyticus* RIMD2210633.

## DISCUSSION

Many bacterial pathogens hijack numerous cellular processes in infected cells by performing orthogonal post-translational modifications of host proteins (13, 20). One such modification is ADP-ribosylation, a process in which the ribose moiety of NAD+ is covalently attached to host proteins or nucleic acids (22–24, 52). In addition to the canonical ARTs such as Diphtheria, ExoA, and Cholix toxins, among others, during the last decade, an increasing number of bARTTs have been found to be delivered by specialized secretion systems, including the T3SS and T6SSs (29–33, 53, 54). In this study, we provide evidence that the VPA1312 ORF encodes a novel T3SS2 effector, named VopU, which corresponds to a new ART protein of the H-Y-[EDQ] family of bARTTs.

Previous work has linked VopU to the T3SS2, as it has been determined that it is part of the bile-induced regulon of VtrB and that its secretion is altered in the absence of critical components of the secretion apparatus such as the VgpA protein (38, 39), but its role as an effector protein has not been analyzed previously. Translocation experiments showed that delivery of VopU into infected host cells occurred independently of the previously characterized chaperone VocC (**Figure 1**). VocC has been shown to be required for the translocation of VopC during infection and is the only chaperone described for the T3SS2 of *V. parahaemolyticus* so far (44). Therefore, it is possible that either VopU requires an as-yet-uncharacterized chaperone or that it possesses a chaperone-independent mechanism for translocation into host cells. In this context, the relatively small size of VopU and the absence of a conserved chaperone-binding site suggest a chaperone-independent mechanism of translocation. Interestingly, it has recently been shown that VopU secretion is affected by the VgpA protein, and bacterial two-hybrid experiments have shown that VgpA and VopU interact (38). Whether the VopU interaction with VgpA contributes to its interaction with the sorting platform and to efficient translocation into infected host cells remains to be tested.

Sequence and structure-based analyses provided evidence that VopU belongs to a novel family of effector proteins with predicted mono-ART activity, distributed across species beyond the *Vibrionaceae* family. As T3SS effector proteins are often linked to horizontally acquired elements, it is plausible that VopU has been transferred between species through horizontal gene transfer. In this context, VopU is often encoded in the vicinity of T3SS2 gene clusters, flanked by IS elements and genes encoding putative transposases.

An interesting observation was the presence of VopU homologs encoded within plasmids belonging to the pO55 family of *E. coli* O55:H7 strains which also encode for the non-LEE T3SS effector NleA. The O55:H7 serotype is considered the progenitor of the contemporary Shiga toxin-containing *E. coli* (STEC) O157:H7 serotype (55) which harbors a related plasmid (pO157:H7) very similar to pO55 plasmids (56) but it does not encode a VopU homolog. The presence of genes encoding VopU homologs within the plasmids present in strains of the ancestor serotype and its absence in contemporary isolates suggest that the loss of VopU might have given the current STEC strains a competitive advantage. In addition to T3SS, we identified VopU homologs in the vicinity of gene clusters encoding other protein secretion systems. In this sense, the presence of genes encoding VopU homologs near T6SS gene clusters of *Photorhabdus hainanensis* and *Shewanella* species strains is intriguing. Recent reports have highlighted the contribution of non-canonical ARTs to the antibacterial activity of the T6SS in *Pseudomonas aureginosa* (RhsP2), *Salmonella* Typhimurium (Tre^Tu^) (29), *Serratia proteamaculans* (Tre1)(31) and *Photorhabdus laumondii* (Tre23) (34). The presence of genes encoding VopU homologs in these clusters suggests that these proteins may also have been repurposed for other protein secretion systems.

Most bacterial mono-ARTs rely on a conserved H-Y-D or H-Y-E catalytic triad, in which the histidine and tyrosine residues are required for NAD binding, while glutamate or aspartic acid is important for catalysis (22, 23). On the other hand, in the human Poly-(ADP-ribose) polymerase (PARP) family of proteins, there is more variability within the catalytic core motif with additional amino acids involved in catalysis (H-Y-[EIYLV]) often linked to their mono or poly-ART activity (57–59). Notably, both bioinformatic analyses of the VopU structure and functional ADP-ribosylation assays in host cells indicate that VopU harbors a unique catalytic motif consisting of an H-Y-Q catalytic triad. To the best of our knowledge, this is the first description of a naturally occurring mono-ART with an H-Y-Q catalytic triad. Even though there are no reports of naturally occurring H-Y-Q ARTs, experiments have been performed that replaced the catalytic glutamate (E) for glutamine (Q) in bacterial and human H-YE ARTs.

For example, when the H-Y-E triad of Diphtheria toxin was changed to H-Y-Q, it did negatively affected NAD binding, but it partially reduced it ART activity while increasing the toxin NAD glycohydrolase activity (60). On the other hand, experiments performed in human PARP1, a poly-ART, revealed that the change of a glutamate for glutamine (E988Q mutant) in the catalytic triad abolished poly-ART activity while maintaining mono-ART activity (61–63). These data indicate that the glutamine residue is required for poly-but not for mono-ART activity. This conclusion was also supported by the fact that in several human PARPs that act mainly as mono-ARTs, glutamate is replaced by valine, leucine, or isoleucine in the catalytic triad (H-Y-[IYLV]) (57–59).

The presence of mono-ADP-ribosylated protein of approximately 18 kDa in cells expressing VopU, suggests that this effector has mono-ART activity, but it raises the question of whether it could also function as a NAD glycohydrolase within host cells, given the overall cellular stress caused by VopU expression within transfected cells (**Figure 5D**). Further biochemical characterization of VopU is needed to determine the kinetics of both ART and potential NAD glycohydrolase activities, and to assess their implications during *Vibrio* infection.

Analysis of the predicted structure of VopU revealed that the β-sheet core region, important for NAD binding, is conserved compared with other bacterial ARTs, such as ExoA (**Figure 2**), but the overall secondary structures of the two proteins differed. These structural differences suggest that VopU might not target the same host protein as previously described toxins of the H-Y-[EDQ] family of bacterial ARTs. Toxins such as ExoA and Diphtheria ADP-ribosylate the diphtamide residue of eEF2, halting protein synthesis and eventually causing host cell death (26, 27).

Several lines of evidence suggest that VopU does not target eEF2. First, expressing VopU in HeLa cells or infecting HT-29 cells with *V. parahaemolyticus* strains harboring or lacking VopU, identified an ADP-ribosylated protein of approximately 18 kDa, whereas eEF2 is approximately 95 kDa. Secondly, both previous work (37) and our infection and heterologous expression data show that VopU does not contribute to host cell death. Finally, the structural differences and a distinct catalytic triad mentioned above also suggest a potentially different target for VopU.

*In vitro* experiments did not identify a contribution of VopU to either invasion, intracellular survival, or host cell cytotoxicity during infection, even though infection with a strain lacking VopU altered the characteristic morphological changes observed during *V. parahaemolyticus* infection. Nevertheless, it is unclear whether this phenotype is a direct or indirect consequence of VopU ART activity.

Expression of VopU in cells altered cellular morphology and induced gene expression changes associated with cellular stress. It is important to mention that both phenotypes could be a consequence of reduced NAD levels caused by VopU heterologous expression and might not reflect the processes altered by physiological levels of VopU translocated by *Vibrio* during infection. In this context, these phenotypes are more informative regarding the biochemical activity of VopU rather than its contribution to bacterial infection. This is supported by RNA-seq analysis, which showed increased expression of host genes associated with cellular stress. One of the genes with the largest fold change in expression was the mitochondrial citrate carrier SLC25A1, which has been shown to be critical to maintain the NADP+/NADPH ratio within the cell, especially in response to stress (64), suggesting that increased expression of SLC25A1 due to expression of VopU is part of the stress response by reducing NAD levels. Similarly, overexpression of Fibroblast growth factor 21 (FGF21) was also observed. It has been shown that increased levels of FGF21 can lead to an increase in cellular NAD^+^ levels, which subsequently activates NAD+-dependent enzymes (65). In this context, further work is needed to determine the exact contribution of VopU during bacterial infection.

Interestingly, VopU is the second bARTT effector protein encoded within the T3SS2 gene cluster together with VopT. We provide evidence that VopT targets additional, unidentified host proteins beyond the previously identified target Ras (36). This suggests that both VopU and VopT can manipulate host NAD to ADP-ribosylate distinct sets of host proteins, ultimately contributing to *Vibrio* infection. Further work is needed to determine whether both effector proteins act in synergy or on distinct host cell processes during infection, which will expand our current understanding of how *Vibrio* utilizes post-translational modifications of host proteins to successfully infect host cells. Finally, it is important to note that, until recently (50, 51, 66), detection of endogenous ADP-ribosylation has been limited by the lack of well-developed and specific reagents. In this context, we think it is important to highlight that the mono-selective ADP-ribose antibody (AbD43647) used in our work to profile ADP-ribosylated proteins is from a novel set of detection reagents with enhanced specificity, recently made commercially available (50, 51, 66). The AbD43647 antibody clone used in this work is specific for mono-ADPr and has a preference for Ser-mono-ADPr (50, 51) suggesting that VopU ADP-ribosylates its target protein at a serine residue. The availability of these novel tools holds the promise of providing better insight into how bacterial pathogens subvert post-translational modifications, such as ADP-ribosylation, to disrupt host cell processes.

Overall, we have identified and characterized VopU, an effector of a novel family of mono-ART with a unique H-Y-Q catalytic triad. Further work on how VopU and VopT manipulate host NAD to ADP-ribosylate distinct sets of host proteins will shed light on how *Vibrio* uses orthogonal post-translational modifications to subvert host biology.

## MATERIALS AND METHODS

### Bacterial strains and growth conditions

The complete list of bacterial strains and plasmids are listed in **Table S1**. *V. parahaemolyticus* RIMD2210633 (5) and its Δ*vscn1*, Δ*vscn2*, and Δ*vscn1* Δ*vscn2* derivatives (67) were used in this study. Bacterial strains were routinely cultured in LB medium or on LB agar plates with 1% NaCl at 37°C. When appropriate, the culture media was supplemented with 0.04% bovine and ovine bile (Sigma catalog no. B8381); 5LJμg/ml and 20 μg/ml chloramphenicol for *V. parahaemolyticus* and *E. coli* strains respectively; 100 ug/ml ampicillin for *E. coli* strains; 1 mM isopropyl-β-d-thiogalactopyranoside (IPTG) to induce expression vector pCyaA in translocation assays. The *V. parahaemolyticus* VPA1312 (VopU) isogenic mutant strain was constructed via standard allelic exchange using a derivative of the pDM4 suicide vector carrying DNA sequences flanking the VPA1312 ORF targeted for deletion, as described previously (67). Plasmids were propagated in *E. coli* DH5-α, and *E. coli* SM10λPir was used to deliver the pDM4 suicide vector via conjugation (68).

### Eukaryotic cell lines and culture conditions

HEK293T, HeLa (ATCC CCL-2), HT-29 (ATCC HTB-28), and Caco-2 (ATCC HTB-37) cells were routinely cultured and maintained in Dulbecco’s modified Eagle medium (DMEM high glucose) (Gibco 11965092) supplemented with 10% fetal bovine serum (FBS) (Cytiva SV30160.03) (DMEM−15% FBS) at 37°C in 5% CO_2_ atmosphere. Cells were routinely passaged at 70-80% confluence.

### Translocation of effector-CyaA fusion proteins

Translocation of effector proteins was analyzed using a CyaA reporter fusion-based translocation assay, as previously described (69). Briefly, HT-29 or Caco-2 cells were seeded at 1.5LJ×LJ10^4^ cells/well in a 96-well plate and cultured in DMEM supplemented with 10% FBS at 37°C and 5% CO_2_ for 48 hours. *V. parahaemolyticus* RIMD2210633 Δ*vscn1* and Δ*vscn1* Δ*vscn2* strains containing pCyaA, pVopV-CyaA or pVopU-CyaA were grown until they reached an OD_600_ of 0.6 in LB medium supplemented with 0.04% bile. The infection assays were performed for 1 h at 37°C and 5% CO_2_ and at a multiplicity of infection (MOI) of 50. The intracellular cyclic AMP (cAMP) levels in host cells were determined using the Intracellular Arbor Assay Cyclic AMP Direct EIA non-acetylated kit (catalog no. K019-H1). Statistical analysis was performed with GraphPad Prism version 11.0 (GraphPad Software, San Diego, California, USA).

### Cytotoxicity and cell viability assays

For experiments aimed at determining T3SS2-dependent cell death, HT-29 or Caco-2 cells were seeded at 8.0LJ×LJ10^4^ cells/well into six-well plates and grown for 2LJdays in complete media. *V. parahaemolyticus* strains were cultured overnight and the next day diluted 1:100 into LB liquid media containing 0.04% bile (to induce T3SS2 expression) and grown for 2 h until attaining an OD_600_ of 0.6. Cells were infected at an MOI ofLJ10 and incubated at 37°C with 5% CO_2_. LDH activity in the medium was analyzed at different time points using the CytoTox 96® Non-Radioactive Cytotoxicity Assay (Promega cat no. G1780). Cell death was also analyzed using the CellTox™ Green Cytotoxicity Assay (Promega cat no. G8741). For experiments aimed at determining cell viability after transfection experiments, cell viability was determined by trypan blue exclusion (0.4% trypan blue) and counted on a hemocytometer (Neubauer cell chamber). Statistical analysis was performed with GraphPad Prism version 11.0 (GraphPad Software, San Diego, California, USA).

### Gentamicin protection assay

Gentamicin protection assays were performed as described in (70) with minor modifications. Briefly, 7×10^4^ HeLa cells/well were seeded in 24-well plate. On the day of infection, bacteria were subcultured 1:100 in LB medium supplemented with bile salts (0.04% w/v) until reach O.D600 = 0.6, cells were infected using MOI = 10 with the respective *V. parahaemolyticus* strains. Infection was synchronized by 5 min of centrifugation at 10 g and infection was allowed for 2 hours at 37°C and 5% CO2. After 2 h of incubation, samples were washed twice with DPBS and incubated with DMEM supplemented with gentamicin 100 ug/mL. At different time points, samples were washed twice with sterile PBS and incubated at room temperature (RT) 10 min with 0.5% of Triton X-100 0.5%. After incubation, samples were serially diluted and plated on LB agar plates and incubated at 37°C overnight for CFU determination, normalizing to the input CFUs. Statistical analysis was performed with GraphPad Prism version 11.0 (GraphPad Software, San Diego, California, USA).

### Fluorescence labeling of infected cells

HeLa cells were seeded at a density of 5×10^4^ cells/ml on glass coverslips in 24-well plates. The day of infection *V. parahaemolyticus* strains harboring the pGFP plasmid (a modified pON.mCherry plasmid (71) where the mCherry ORF was replaced with the EGFP protein of plasmid pFCcGi (72)) were grown in LB medium supplemented with 0.04% bile until they reached an OD_600nm_ of 0.6. Cells were infected at a MOI=10 and at each time point cells were washed with sterile PBS and fixed with 4% PFA for 10 minutes. Fixed cells were washed three times and incubated with permeabiization/blocking solution (5% BSA, 0.1% saponin). Coverslips were incubated for 1 hour with 1:1000 Hoechst and 1:100 Alexa 467 phalloidin solution followed by three washes with blocking buffer. Coverslips were mounted on glass slides using ProLong Diamond antifade mounting media. Images were acquired using confocal microscope Spinning Disk (Olympus IXplore SpinSR). Analysis of Z-stack, orthogonal planes and 3D reconstruction was performed using Imaris 10.2 software and ImageJ 1.54j.

### Heterologous expression of VopU within eukaryotic cells

A codon optimized version of VopU with a C-terminal HA epitope was obtained from Genewiz (USA). VopU variants with mutations in the H-Y-Q catalytic triad were generated with the Q5® Site-Directed Mutagenesis kit (NEB E0554S) using divergent primers. Primers were designed using the SnapGene software (from Dotmatics; available at snapgene.com). Each gene was cloned in plasmid pAcGFP1-N1 (Clonetech Takara), replacing the AcGFP1 ORF using Gibson Assembly (73), allowing gene expression under the constitutive CMV promoter. Whole plasmid sequencing for each construct was performed by Plasmidsaurus using Oxford Nanopore Technology with custom analysis and annotation. Transfection experiments were performed using the TransIT-LT1 transfection reagent (Mirus Bio Cat No. MIR 2300) following the manufactureŕs recommendations.

### RNA-seq analysis

For RNA-sequencing experiments of transfected cells, 10×10^5^ cells per condition, and in duplicate, were collected at 48 hours post transfection by washing once with DPBS and dissociated with TrypLE Express (Gibco), collected by centrifugation and resuspended/lysed in 50 µl DNA/RNA Shield (Zymo R1100-50). RNA-Seq was performed by Plasmidsaurus using Illumina Sequencing Technology with the in-house custom analysis and annotation pipeline.

### ADP-ribosylation analysis and Immunoblotting

Whole cell lysates from transfected and non-transfeced HeLa cells were obtained from 50×10^4^ cells treated with Pierce IP Lysis Buffer (Cat no. 87787) supplemented with 250 U/ul of Benzonase nuclease (Cat no. 71205-3) and 1X Halt Protease and Phosphatase Inhibitor Cocktail EDTA-free (Cat no. 78441). Lysis was performed in ice for 5 minutes. The cell lysate was clarified by centrifugation at 13.000 rpm for 10 minutes at 4°C. For whole cell lysates of HeLa cells infected with *V. parahaemolyticus* strains, an additional filtration step with a 0.22um PVDF filter was performed after clarification of the cell lysate to remove any potential bacterial contaminant. Protein samples were prepared in 1X Laemli buffer and heated for 1 minute at 60°C to prevent ADP-ribosylation loss (74) and loaded onto 12% SDS-PAGE gels. For immunoblot analysis, gels were transferred onto PVDF membranes (Immobilon-P, Cat no. IPVH0010) at 30V for 16 hours at 5°C using a Trans-Blot Turbo System (BioRad) and blocked for 1 hour at RT with blocking solution (5% w/v milk diluted in TBS supplemented with 0.1% Tween). Immunoblots were performed with antisera directed against HA tag (α-HA, 1:1000 dilution, Cat no. ab9110, Abcam), GADPH (14C10) (α-GADPH; 1:1000 dilution; Cat no. 2118, Cell Signaling Technologies). To detect ADP-ribosylation, the Mono-ADP-Ribose antibody AbD43647 (Cat no. TZA020, BioRad) was coupled with Rabbit IgG-FcSpyCatcher3 (Cat no. TZC013, BioRad) following the manufactureŕs recommendations and used at a 1:250 dilution. Secondary antibodies conjugated to horseradish peroxidase and Supersignal West Pico chemiluminescent substrate (Thermo Scientific) were used to detect protein levels, with imaging performed on a C-DiGit Blot Scanner (LI-COR).

### Sequence, structural and phylogenetic analysis

The identification of conserved functional domains and motifs within VopU was performed by analysis of the VPA1312 aminoacid sequence with the InterProScan database (75, 76). Identification of VopU orthologs was carried out using the VPA1312 amino acid and nucleotide sequences as queries in BLASTp, BLASTn, BLASTx, tBLASTn, and tBLASTx analyses (77) using publicly available bacterial genome sequences of the NCBI database (January 2026). A 94% sequence length, 70% identity and 80% sequence coverage threshold were used to select positive matches. The list of identified homologs is part of **Table S2**. Sequence conservation was analyzed by multiple sequence alignments T-Coffee Expresso (78) and visualized by ESPript 3.0 (79). Comparative genomic analysis was performed using the multiple aligner Mauve (80) and EasyFig v2.2.5 (81). Multiple sequence alignments were used for phylogenetic analyses that were performed with the Molecular Evolutionary Genetics Analysis (MEGA) software version 12.0 (82) and visualized by iTOL (83). Phylogenetic trees were built from the alignments by the bootstrap test of phylogeny (2000 replications) using the maximum-likelihood (ML) method with a Jones-Taylor-Thornton (JTT) correction model. For protein structure prediction, AlphaFold3 (84) was used. Protein structure visualization, and superposition were performed using Mathmaker of UCSF ChimeraX (85). Protein structure searchers were performed with the Foldseek server (86).

## ACKNOWLEDGMENTS

We thank members of the Blondel Laboratory for helpful discussions on all aspects of this project and for their comments on the manuscript. We thank Cristobal Blondel and Carmen Blondel for technical assistance.

## CONFLICT OF INTERESTS

The authors declare that there are no conflicts of interest.

## FUNDING INFORMATION

This work was funded by the Howard Hughes Medical Institute (HHMI)-Gulbenkian International Research Scholar Grant #55008749 and FONDECYT Grant 1241637 (ANID). CS was supported by FONDECYT grant 1212075. SA was supported by ANID grant 21210879.

## Notes

### Competing Interest Statement

The authors have declared no competing interest.

